# The Inflammasome Pathway is Activated by Dengue Virus Non-structural Protein 1 and is Protective During Dengue Virus Infection

**DOI:** 10.1101/2023.09.21.558875

**Authors:** Marcus P. Wong, Evan Y.W. Juan, Sai S. Chelluri, Phoebe Wang, Felix Pahmeier, Bryan Castillo-Rojas, Sophie F. Blanc, Scott B. Biering, Russell E. Vance, Eva Harris

## Abstract

Dengue virus (DENV) is a medically important flavivirus causing an estimated 50-100 million dengue cases annually, some of whom progress to severe disease. DENV non-structural protein 1 (NS1) is secreted from infected cells and has been implicated as a major driver of dengue pathogenesis by inducing endothelial barrier dysfunction. However, less is known about how DENV NS1 interacts with immune cells and what role these interactions play. Here we report that DENV NS1 can trigger activation of inflammasomes, a family of cytosolic innate immune sensors that respond to infectious and noxious stimuli, in mouse and human macrophages. DENV NS1 induces the release of IL-1β in a caspase-1 dependent manner. Additionally, we find that DENV NS1-induced inflammasome activation is independent of the NLRP3, Pyrin, and AIM2 inflammasome pathways, but requires CD14. Intriguingly, DENV NS1-induced inflammasome activation does not induce pyroptosis and rapid cell death; instead, macrophages maintain cellular viability while releasing IL-1β. Lastly, we show that caspase-1/11-deficient, but not NLRP3-deficient, mice are more susceptible to lethal DENV infection. Together, these results indicate that the inflammasome pathway acts as a sensor of DENV NS1 and plays a protective role during infection.

## Introduction

Dengue virus (DENV) is a mosquito-borne flavivirus consisting of 4 serotypes (DENV1-4) that represents a growing burden on global public health, with cases increasing 10-fold over the past 20 years. Over 3.8 billion people are at risk of infection with DENV, estimated to reach 6.1 billion by 2080 as urban populations grow and climate change increases the suitable range for *Aedes* mosquitoes, the transmission vectors for DENV.^1^ Of the estimated 105 million people infected by DENV annually, up to 51 million develop dengue; symptoms span a wide range of clinical outcomes from an acute febrile illness accompanied by joint and muscle pain to severe disease characterized by vascular leakage and thrombocytopenia, hemorrhagic manifestations, pulmonary edema, and hypovolemic shock.^2, 3^ The causes of endothelial dysfunction and vascular leak seen in severe dengue disease are likely multifactorial, but some studies suggest a “cytokine storm” triggered by uncontrolled viral replication and immune activation.^4, 5^ There are no current treatment options for severe dengue disease other than supportive care, due in part to an incomplete understanding of dengue pathogenesis.^6^ This underscores a need to better understand both protective and pathogenic host pathways to develop future therapeutics for dengue.

DENV non-structural protein 1 (NS1) has drawn recent interest as a vaccine and therapeutic target for the prevention of severe dengue. DENV NS1 is an approximately 45-kDa protein that dimerizes after translation in infected cells.^7^ Dimeric, intracellular NS1 associates with the lumen of the endoplasmic reticulum and participates in the formation of the viral replication complex.^9–, 11^ NS1 is also secreted from infected cells as a tetramer and/or hexamer, with NS1 dimers oligomerizing around a lipid cargo enriched in triglycerides, cholesteryl esters and phospholipids.^7, 11, 12^ Secreted NS1 plays multiple roles during infection, including binding and inactivating complement components and interacting directly with endothelial cells to induce endothelial hyperpermeability and pathogenic vascular leak.^13–17^ Mice vaccinated with DENV NS1 are protected from lethal systemic DENV challenge in mouse models of infection, and blockade of NS1-induced endothelial hyperpermeability by glycan therapeutics or by NS1-specific monoclonal antibodies also reduces DENV-induced disease, emphasizing the pivotal roles NS1 plays in DENV replication and pathogenesis.^14, 18–21^ DENV NS1 has also been shown to induce the activation of pro-inflammatory cytokines such as TNF-α and IL-6 in both murine and human macrophages.^22–25^ Studies in mice have identified a TLR4-dependent axis for pro-inflammatory cytokine induction, and it has been hypothesized that NS1-induced macrophage activation leads to cytokine storm and immunopathology; however, few studies have experimentally assessed the mechanisms by which NS1 induces inflammation and whether NS1-induced inflammation contributes to overall viral pathogenesis.

DENV has been shown to trigger multiple innate immune pathways that contribute to both host defense and pathogenesis.^26, 27^ Among these pathways are the inflammasomes, a class of innate immune sensors that surveil the cytosol for a broad range of pathogen or damage-associated molecular patterns (PAMPs/DAMPs).^28^ Canonical inflammasomes recruit the cysteine protease caspase-1 via the apoptosis-associated speck-like protein containing CARD (ASC) protein.^29^ Certain inflammasomes respond to PAMPs such as viral double-stranded DNA in the case of the AIM2 inflammasome, or can be triggered by pathogenic effectors; examples include sensing of viral protease activity by the NLRP1B and CARD8 inflammasomes, sensing of ion fluxes and membrane damage by the NLRP3 inflammasome, or sensing of toxin-induced Rho guanosine triphosphatase (Rho GTPase) inactivation by the pyrin inflammasome.^28, 30–33^ Further, caspase-11 in mice and caspases-4 and -5 in humans can activate the non-canonical inflammasome, in which caspase-11/4/5 binding to lipid A from bacterial lipopolysaccharide (LPS) leads to activation of the NLRP3 inflammasome.^34, 35^

Inflammasome signaling typically comprises a two-step process in which inflammasome components and substrates are first transcriptionally upregulated and/or ‘primed’, usually in response to PAMPs/DAMPs and nuclear factor-κB (NF-κB) signaling.^28^ After priming, a second stimulus induces inflammasome oligomerization, leading to ASC recruitment and caspase-1 autoproteolytic processing into its active form.^29^ The active caspase-1 protease can then cleave pro-IL-1β, pro-IL-18 and gasdermin D (GSDMD) into their bioactive forms. Cleavage of GSDMD leads to insertion and oligomerization of the N-terminal domain (GSDMD-NT) to form pores in the plasma membrane.^36^ The formation of GSDMD pores canonically leads to pyroptosis, a form of inflammatory cell death; however, recent work has shown that GSDMD pore formation and pyroptosis are distinct events and that macrophages can release IL-1β from GSDMD pores without undergoing pyroptosis in response to certain stimuli.^37–41^ GSDMD pores also facilitate the release of cleaved IL-1β and IL-18, which then serve as major mediators of inflammation contributing to host defense as well as driving immunopathology.^37, 42^ Many viruses have been shown to activate inflammasomes during infection, including influenza A virus, HIV, SARS-CoV-2, picornaviruses, and DENV.^27, 33, 43–45^ Inflammasome activation by viruses can be protective and/or can contribute to pathogenic outcomes.^43, 45–48^ Ultimately, the impact of inflammasome activation depends on the context and timing of the infection; thus, understanding these complexities is crucial for designing inflammasome-targeted therapies as potential treatments for viral disease.

Several studies have begun to investigate whether DENV infection triggers inflammasome activation and how this might impact DENV pathogenesis. Studies have shown that *in vitro* DENV infection of mouse and human macrophages, human skin endothelial cells, and platelets, as well as infection in mice can induce inflammasome activation.^49–54^ Clinically, IL-1β levels are also elevated in dengue patients, implicating a role for inflammasome activation in human DENV infections.^53, 55^ Mechanistic studies have implicated both the membrane (M) and NS2A/B proteins of DENV as viral triggers of the NLRP3 inflammasome.^49, 52^ *In vivo* studies using an adeno-associated virus (AAV) vector to induce DENV M expression suggested that DENV M can cause NLRP3-dependent vascular leak, though the relevance of M-induced inflammasome activation in DENV infection is unknown.^49^ Another study showed that mice treated with an IL-1 receptor antagonist during DENV infection lost less weight and experienced less vascular leak compared to untreated controls.^53^ Although it is established that DENV infection can induce inflammasome activation, the viral triggers and the contribution of inflammasome activation during infection remain open areas of investigation. In this study, we identify secreted NS1 as a trigger of the inflammasome pathway in a caspase-1-dependent manner that is independent of the NLRP3, pyrin, and AIM2 inflammasomes but dependent on CD14. Further, we demonstrate that caspase-1/11 deficiency, but not NLRP3 deficiency, makes mice more susceptible to DENV infection, indicating that inflammasome activation can be protective during DENV infection.

## Results

### DENV NS1 can activate the inflammasome pathway

Since DENV NS1 is secreted from infected cells and can activate macrophages to induce a pro-inflammatory response, we hypothesized that DENV NS1 could be a viral trigger of the inflammasome pathway in macrophages.^22^ To test this hypothesis, we assessed whether DENV NS1 could activate the inflammasome pathway in mouse bone marrow-derived macrophages (BMDMs). BMDMs were first primed with PAM_3_CSK_4_, a synthetic triacylated lipopeptide TLR1/TLR2 agonist, and then treated with DENV NS1. Cell supernatants were assessed 24 hours (h) post-treatment for the presence of IL-1β as a readout of inflammasome activation. We found that DENV NS1 induced the release of IL-1β in BMDMs in a dose-dependent manner **(Figure 1A)**. Additionally, DENV NS1 induced the cleavage of pro-IL-1β, GSDMD, and pro-caspase-1 into their cleaved, bioactive components (IL-1β p17, GSDMD-N p31, and caspase-1 p20, respectively), confirming activation of the inflammasome pathway **(Figure 1B)**. Next, we obtained BMDMs from mice genetically deficient in caspase-1 and -11, required for canonical and non-canonical inflammasome signaling, respectively, and compared them to BMDMs from WT caspase-1/11-sufficient littermates and found that NS1-induced IL-1β release was abolished in BMDMs derived from caspase-1/11 knockout mice **(Figure 1C)**. Similarly, nigericin, a NLRP3 inflammasome agonist, was unable to induce IL-1β release in caspase-1/1-deficient macrophages **(Figure 1C)**. Consistent with these findings, the caspase-1-specific inhibitor AC-YVAD-cmk inhibited both DENV NS1 and nigericin-induced IL-1β release in a dose-dependent manner (**Figure 1D).**^56^ Additionally, DENV NS1 was also able to induce cleavage of caspase-1 and the release of bioactive IL-1β in human THP-1-derived macrophages in a caspase-1 dependent manner **(Figure 1E-F).** Collectively, these data suggest that DENV NS1 induces inflammasome activation in a caspase-1-dependent manner in macrophages.

**Figure 1.**
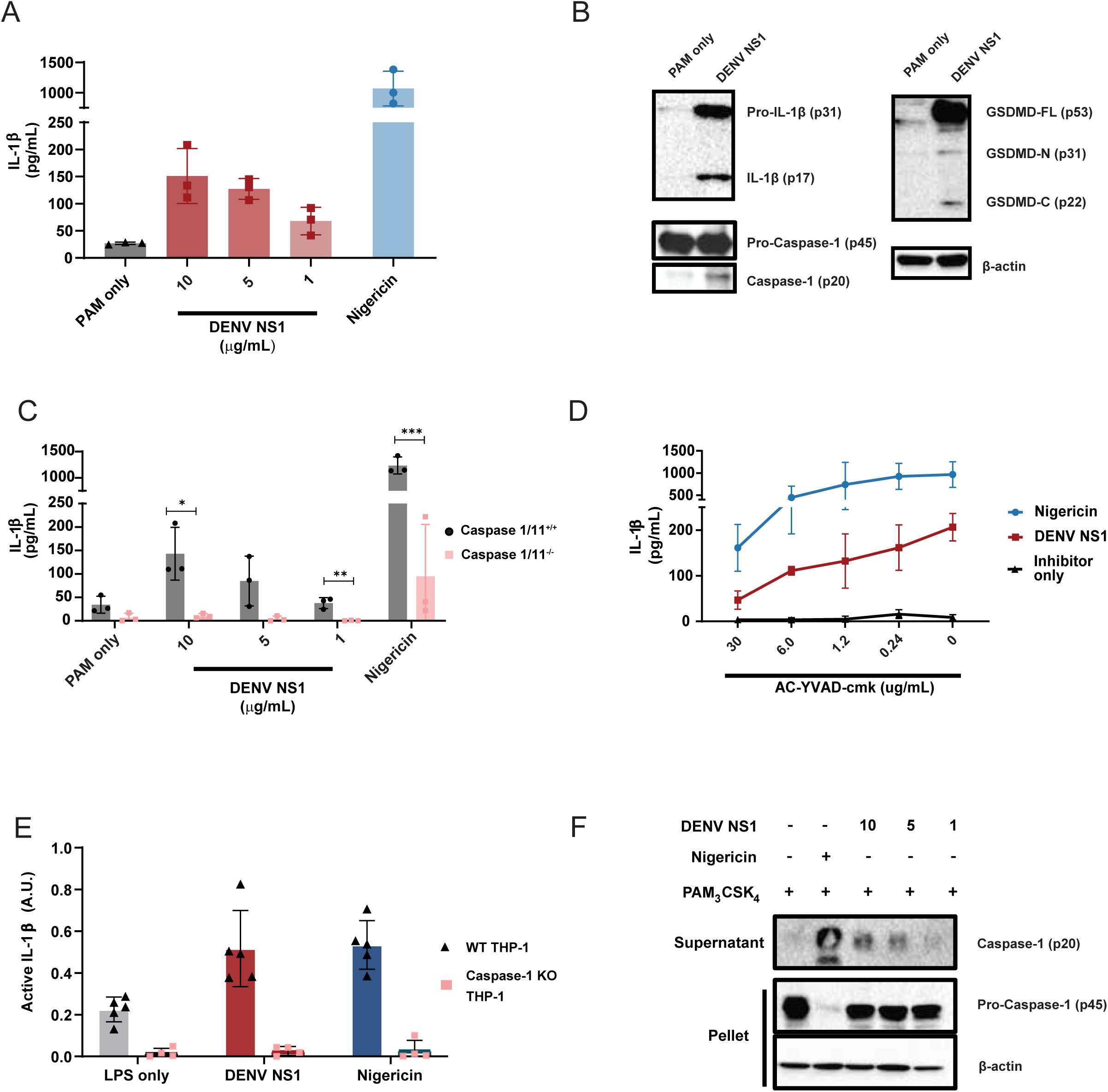
DENV NS1 can activate the inflammasome. **(A)** BMDMs were primed with PAM_3_CSK_4_ (1μg/mL) for 17 h and then treated with DENV2 NS1 at indicated concentrations, treated with 5μM nigericin, or left untreated (PAM only). IL-1β levels in the supernatant after 2h (nigericin) or 24h (NS1 and PAM only) were measured by ELISA. *p<0.05 as determined by one-way ANOVA with Dunn’s multiple comparison correction. **(B)** Representative Western blots of BMDM cell lysates after priming with PAM_3_CSK_4_ (1μg/mL) for 17h and treatment with 10ug/mL DENV2 NS1 (DENV NS1) or PAM_3_CSK_4_ treatment for 24h without NS1 treatment (PAM only). **(C)** WT and *Casp1/11^-/-^* BMDMs were primed with PAM_3_CSK_4_ (1μg/mL) for 17h and then treated with DENV2 NS1 at the indicated concentrations, nigericin (5μM), or medium (PAM only). IL-1β levels in the supernatant after 2h (Nigericin) or 24h (NS1 and PAM only) were measured by ELISA. *p<0.05, **p<0.01. Statistical significance was determined using two-way ANOVA followed by multiple t-tests using Holm-Sidak correction. **(D)** BMDMs were primed with PAM_3_CSK_4_ (1μg/mL) for 17h and then pre-treated with Ac-YVAD-cmk at the indicated concentrations before addition of DENV2 NS1 (10μg/mL), nigericin (5μM), or medium (Inhibitor only). IL-1β levels in supernatant 2h (Nigericin) or 24h (NS1 and PAM only) were measured by ELISA. **(E)** WT or *Casp-1^-/-^*THP-1 human monocytes were differentiated into macrophages in 10ng/mL PMA and primed with medium or LPS for 4h. Primed macrophages were treated with DENV NS1 (10μg/mL) or left untreated (LPS only). Eighteen hours later, supernatants were collected. Cells were stimulated with 5μM nigericin for 2h as a positive control. Bioactive IL-1β in supernatants was measured using HEK-Blue IL-1R reporter cells. **(F)** Representative Western blots of THP-1 macrophage cell lysates after priming with PAM_3_CSK_4_ (100ng/mL) for 17h and treatment with DENV2 NS1 at indicated concentrations (μg/mL), treatment with 5μM nigericin, or no treatment for 24h. The data are shown as the mean ± standard deviation (SD) of 3 independent experiments (A, C-D) or 4-6 independent experiments (E) or a representative image taken from 2 biological replicates (B,F).

### DENV NS1-mediated inflammasome activation is NLRP3-independent

Since the NLRP3 inflammasome has previously been shown to be activated by other DENV proteins as well as by NS1 from the closely related Zika virus, we hypothesized that DENV NS1 might also activate the NLRP3 inflammasome. ^49, 52, 57^ Surprisingly, we found that DENV NS1 was still able to induce IL-1β cleavage and release in BMDMs derived from *Nlrp3*^-/-^ mice, whereas IL-1β release was severely reduced in BMDMs derived from *Nlrp3*^-/-^ mice treated with the NLRP3 inflammasome activator nigericin (**Figure 2A-B)**. Likewise, treatment of WT BMDMs with the NLRP3-specific inhibitor MCC950 inhibited nigericin-mediated IL-1β release in a dose-dependent manner, whereas DENV NS1-induced IL-1β release was unaffected by MCC950 treatment **(Figure 2C).**^58^ These data suggest that DENV NS1-mediated inflammasome activation is independent of the NLRP3 inflammasome. Since IL-1β release downstream of the non-canonical inflammasome and the lipopolysaccharide (LPS)-triggered inflammasome pathway requires NLRP3, these results further suggest that NS1-induced inflammasome activation is not via caspase-11 or due to LPS contamination.^59^ We then utilized CRISPR-Cas9 gene editing to knock out additional components in the inflammasome pathway via nucleofection of Cas9-guide RNA (gRNA) ribonucleoprotein complexes in WT BMDMs to attempt to identify the inflammasome pathway activated by DENV NS1. We targeted *Nlrp3* (encoding Nlrp3), *Aim2* (encoding Aim2), *Mefv* (encoding Pyrin) and *Casp1* in C57BL/6 mice using 2 guide RNAs per gene and used a non-targeting guide (NTG) as an experimental control. This approach achieved robust deletion of the target proteins, as assessed by Western blot (**Figure 2D).** We then treated gene-edited BMDMs with DENV NS1 for 48h and assessed inflammasome activation by IL-1β release, as measured by ELISA. Corroborating the results from *Nlrp3^-/-^* mice, CRISPR-Cas9 knockout of *Nlrp3* did not affect DENV NS1-mediated inflammasome activation **(Figure 2E).** CRISPR-Cas9 knockout of the AIM2 and Pyrin inflammasomes also had no effect on DENV NS1-induced inflammasome activation. Thus, DENV NS1-mediated inflammasome activation in BMDMs is independent of the NLRP3, AIM2, and Pyrin inflammasomes.

**Figure 2.**
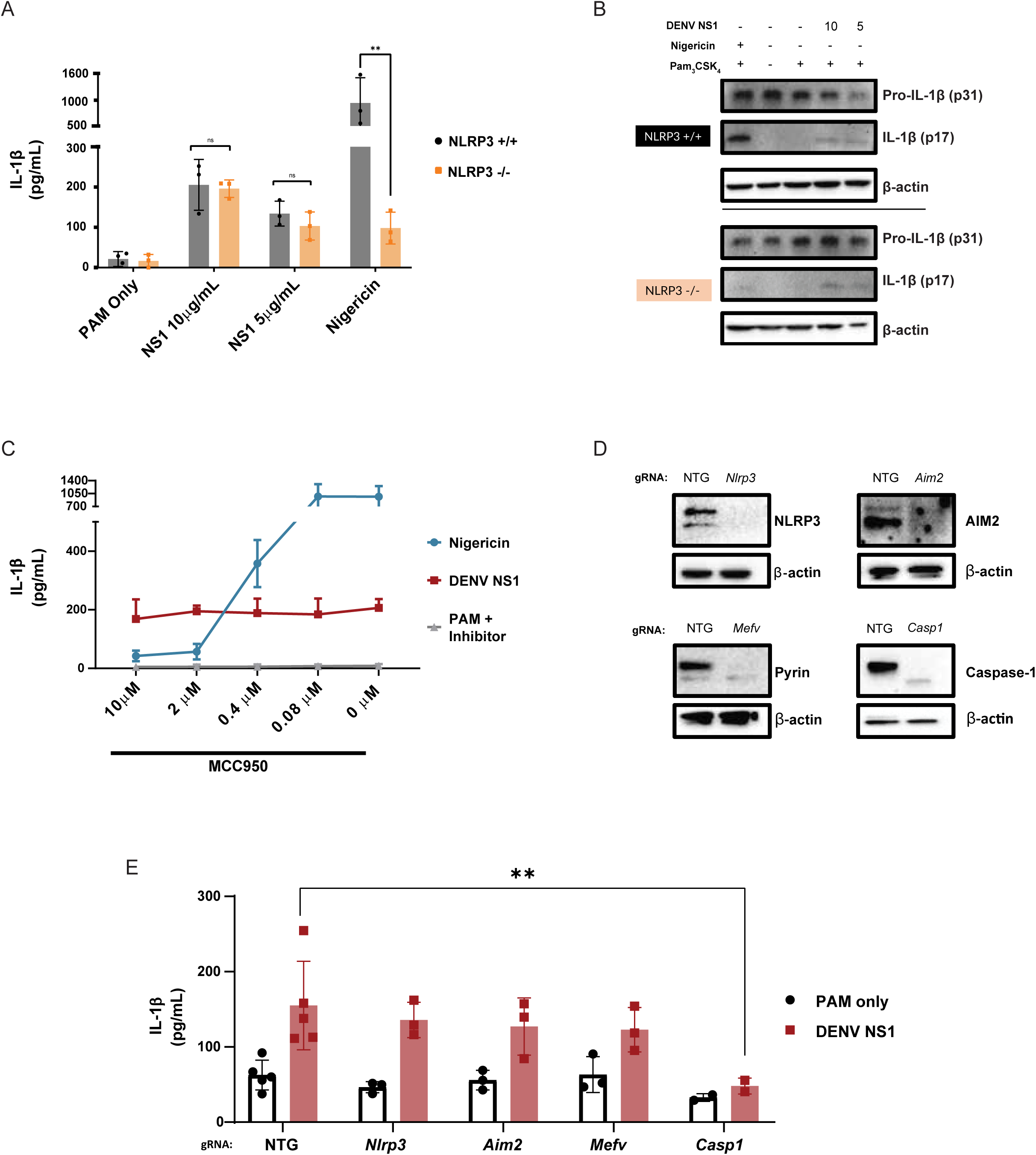
DENV NS1-induced inflammasome activation is NLPR3-independent. **(A)** WT and *Nlrp3 ^-/-^* BMDMs were primed with PAM_3_CSK_4_ (1μg/mL) for 17h and then treated with DENV2 NS1 at indicated concentrations, nigericin (5μM), or medium (PAM only). IL-1β levels in supernatant 2h (nigericin) or 24h (NS1 and PAM only) were measured by ELISA. *p<0.05, **p<0.01. Statistical significance was determined using two-way ANOVA followed by multiple t-tests with Holm-Sidak correction. **(B)** Representative Western blots of cell lysates from WT and *Nlrp3^-/-^* BMDMs after priming with PAM_3_CSK_4_ (1μg/mL) for 17h and treatment with DENV2 NS1 (10 or 5 μg/mL), treatment with nigericin (5μM), or no treatment for 24h. **(C)** BMDMs were primed with PAM_3_CSK_4_ (1μg/mL) for 17h and then pre-treated with MCC950 at the indicated concentrations before addition of DENV2 NS1 (10μg/mL), nigericin (5μM), or medium (Inhibitor only). IL-1β levels in the supernatant after 2h (Nigericin) or 24h (NS1 and PAM only) were measured by ELISA. **(D)** Representative Western blots of cell lysates from BMDMs nucleofected with Cas9-gRNA ribonuclear protein complexes to knock out the indicated genes. Two gRNAs per gene were used per nucleofection. NTG = non-targeting guide. (**E)** Knockout BMDMs from **(D)** were primed with PAM_3_CSK_4_ (1μg/mL) for 17h and treated with DENV2 NS1 (10μg/mL) or left untreated for 48h. *p<0.05. Statistical significance was determined using two-way ANOVA followed by multiple t-tests with Holm-Sidak correction. The data are shown as the mean ± SD of 3 biological replicates (A,C), a representative image taken from 2 biological replicates (B,D), or data pooled from 5 independent experiments with 3 biological replicates per guide (E).

### DENV NS1-induced inflammasome activation is dependent on CD14 and does not lead to cell death

We observed that DENV NS1-treated BMDMs maintain their morphology and do not undergo detectable cell death, in contrast to nigericin-treated cells, despite evidence of cleavage of GSDMD (**Figure 1B)**. Measurement of lactase dehydrogenase (LDH) is often used as a proxy for pyroptotic cell death; consistent with this, we found that PAM_3_CSK_4-_primed BMDMs treated with nigericin rapidly released near maximum amounts of LDH 2h post-treatment.^60–62^ In contrast, PAM_3_CSK_4_-primed, NS1-treated BMDMs released LDH at similar levels to the BMDMs primed with PAM_3_CSK_4_ alone, up to 24h post-treatment **(Figure 3A).** In addition, at 24h post-treatment, nigericin-treated macrophages displayed high levels of staining by an amine-reactive dye used to fluorescently label dead cells, whereas DENV NS1-treated macrophages were labeled at similar levels to the untreated controls **(Figure 3B)**. These lines of evidence indicate that DENV NS1 induces inflammasome activation without inducing cell death. Previous studies have shown that pyroptosis and inflammasome activation are separable processes and that myeloid cells can release IL-1β over time without undergoing pyroptosis.^38, 39, 41^ One such study showed that oxidized phospholipids can enhance IL-1β release without inducing cell death through engagement of CD14 in macrophages and dendritic cells.^63^ Since DENV NS1 is secreted from infected cells as an oligomer surrounding a lipid cargo, we hypothesized that DENV NS1 might also enhance IL-1β release by delivering lipids to cells in a CD14-dependent manner.

**Figure 3.**
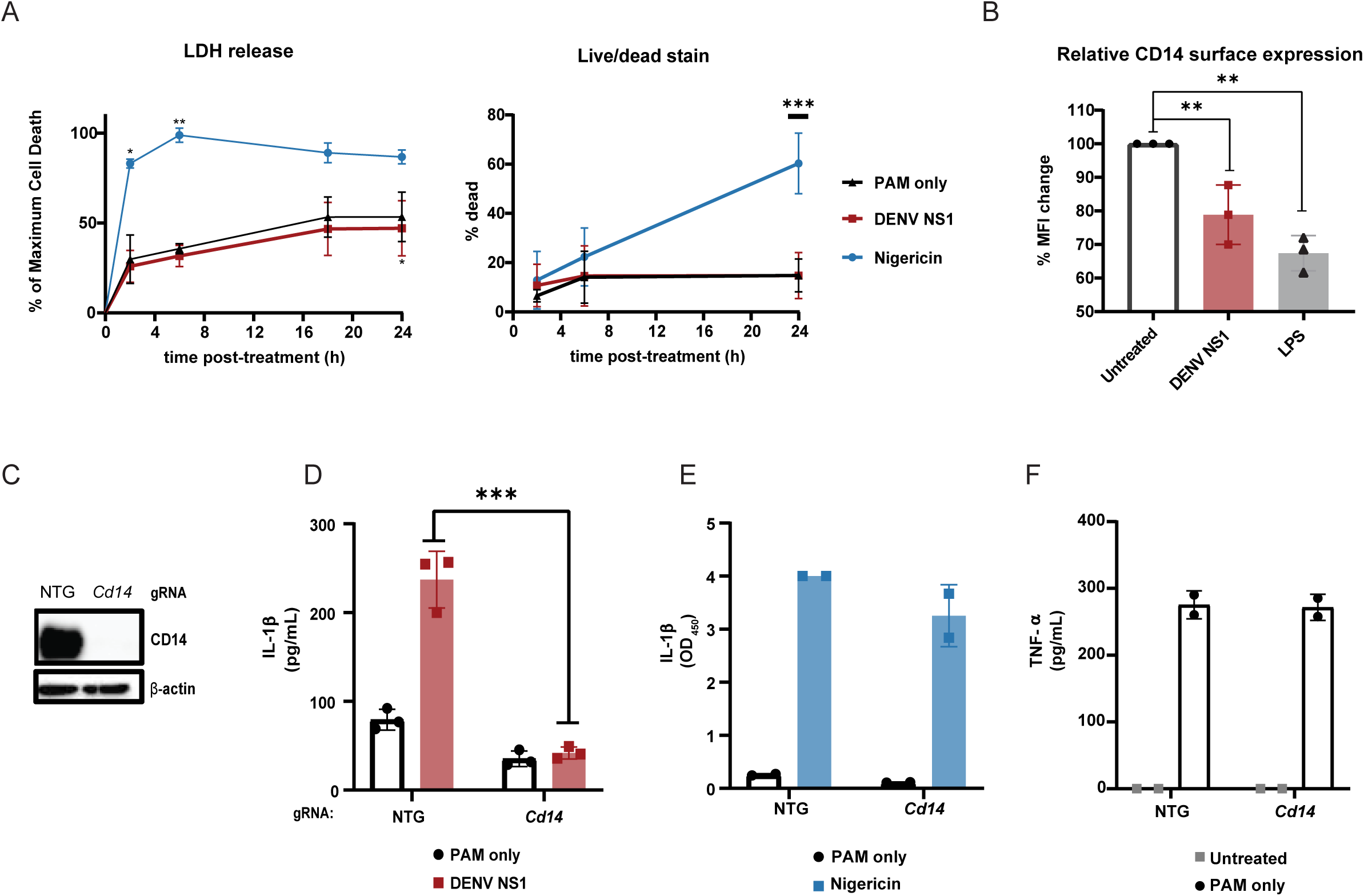
DENV NS1 induces inflammasome activation in macrophages in a CD14-dependent manner without inducing cell death. **(A-B)** BMDMs were primed with PAM_3_CSK_4_ (1μg/mL) for 17h and then treated with DENV2 NS1 (10μg/mL), nigericin (5μM), or medium (PAM only). **(A)** At the indicated timepoints, supernatants were assessed for lactase dehydrogenase (LDH) levels as a proxy for cell death. LDH levels were calculated as a percentage of maximum LDH release. **(B)** Cells were stained using a LIVE/DEAD Fixable Far Red stain and analyzed by flow cytometry. *p<0.05 **p<0.01 ***p<0.001. Statistical significance was determined using two-way ANOVA with Dunnetts’s multiple comparison test. **(C)** BMDMs were primed with PAM_3_CSK_4_ (1μg/mL) for 17h and then treated with DENV2 NS1 (10μg/mL), LPS (5μg/mL) or no treatment (Untreated). After 30 min, cells were stained for surface CD14 expression and analyzed by flow cytometry. Data are normalized as a percentage of the median fluorescence intensity of the treatment groups divided by the untreated control. **p<0.01. Statistical significance was determined using one-way ANOVA with Holm-Sidak’s multiple comparisons test. **(D)** Representative Western blots of cell lysates from BMDMs nucleofected with either NTG or CD14 Cas9-gRNA ribonuclear protein complexes. **(E-G)** BMDMs from **(D)** were primed with PAM_3_CSK_4_ (1μg/mL) for 17h and treated with DENV2 NS1 (10μg/mL), nigericin (5μM) or no treatment for 48h. IL-1β levels in supernatant after 24h (Nigericin) or 48h (NS1 and PAM only) were measured by ELISA **(E-F)**. TNF-α levels were measured in supernatants 17h post-priming with PAM_3_CSK_4_ **(G).** ***p<0.001. Statistical significance was determined using two-way ANOVA with Sidak’s multiple comparison test. The data are shown as the mean ± SD of 3 biological replicates (A,C,E), 5 biological replicates (B), or 2 biological replicates (F-G) or a representative image taken from 2 biological replicates (D).

Indeed, we found that DENV NS1 was able to deplete surface levels of CD14 on BMDMs, as was LPS, a canonical CD14 ligand, suggesting that DENV NS1 can induce endocytosis of CD14 similar to other characterized CD14 ligands **(Figure 3C).** We next generated CD14-deficient BMDMs by nucleofection **(Figure 3D)** and treated these cells with DENV NS1. We found that IL-1β release was abrogated in DENV NS1-treated CD14-deficient BMDMs compared to NTG control cells, suggesting that CD14 is required for DENV NS1 inflammasome activation **(Figure 3E)**. Nigericin-treated CD14 knockout BMDMs still induced IL-1β release at similar levels to NTG controls **(Figure 3F),** and CD14-deficient BMDMs were able to secrete TNF-α in response to PAM_3_CSK_4_ **(Figure 3G)**, showing that NF-κB responses needed for inflammasome priming were intact, leading us to conclude that the lack of NS1-triggered inflammasome activation in CD14-deficient macrophages was not a consequence of off-target effects in other inflammasome pathways. These findings suggest that DENV NS1 inflammasome activation is CD14-dependent and induces Il-1β release without pyroptosis.

### Caspase-1/11 mediates a protective response during DENV infection

We have previously characterized a lethal mouse model of DENV infection and disease consisting of interferon α/β receptor-deficient (*Ifnar ^-/-^*) mice infected with a mouse-adapted DENV2 strain (D220).^64^ To determine how inflammasome activation impacts DENV pathogenesis upon viral infection, we crossed *Casp1/11 ^-/-^* or *Nlrp3 ^-/-^* separately with *Ifnar ^-/-^* mice to generate *Casp1/11 ^-/-^x Ifnar ^-/-^* and *Nlrp3 ^-/-^ x Ifnar ^-/-^*double-deficient mice and infected them with DENV2 D220. We found that *Casp1/11 ^-/-^x Ifnar ^-/-^* mice were significantly more susceptible to lethal DENV2 infection and showed greater morbidity compared to littermate controls with functional *Casp1/11* alleles **(Figure 4A-B)**, indicating that inflammasome activation plays a protective role during DENV2 infection in mice. Previous studies have implicated the NLRP3 inflammasome as playing a pathogenic role during DENV infection; thus, mice deficient in NLRP3 would be predicted to exhibit less severe disease compared to NLRP3-functional mice. However, we found that *Nlrp3 ^-/-^x Ifnar ^-/-^* mice displayed similar outcomes after lethal DENV2 challenge as littermate controls with functional *Nlrp3* alleles across both high and low doses of DENV2 D220 **(Figure 4C-D)**. Thus, these data suggest that while inflammasomes can play a protective role during DENV infection, this protection is independent of the NLRP3 inflammasome, consistent with NS1-triggered inflammasome activation patterns that we measured *in vitro*.

**Figure 4.**
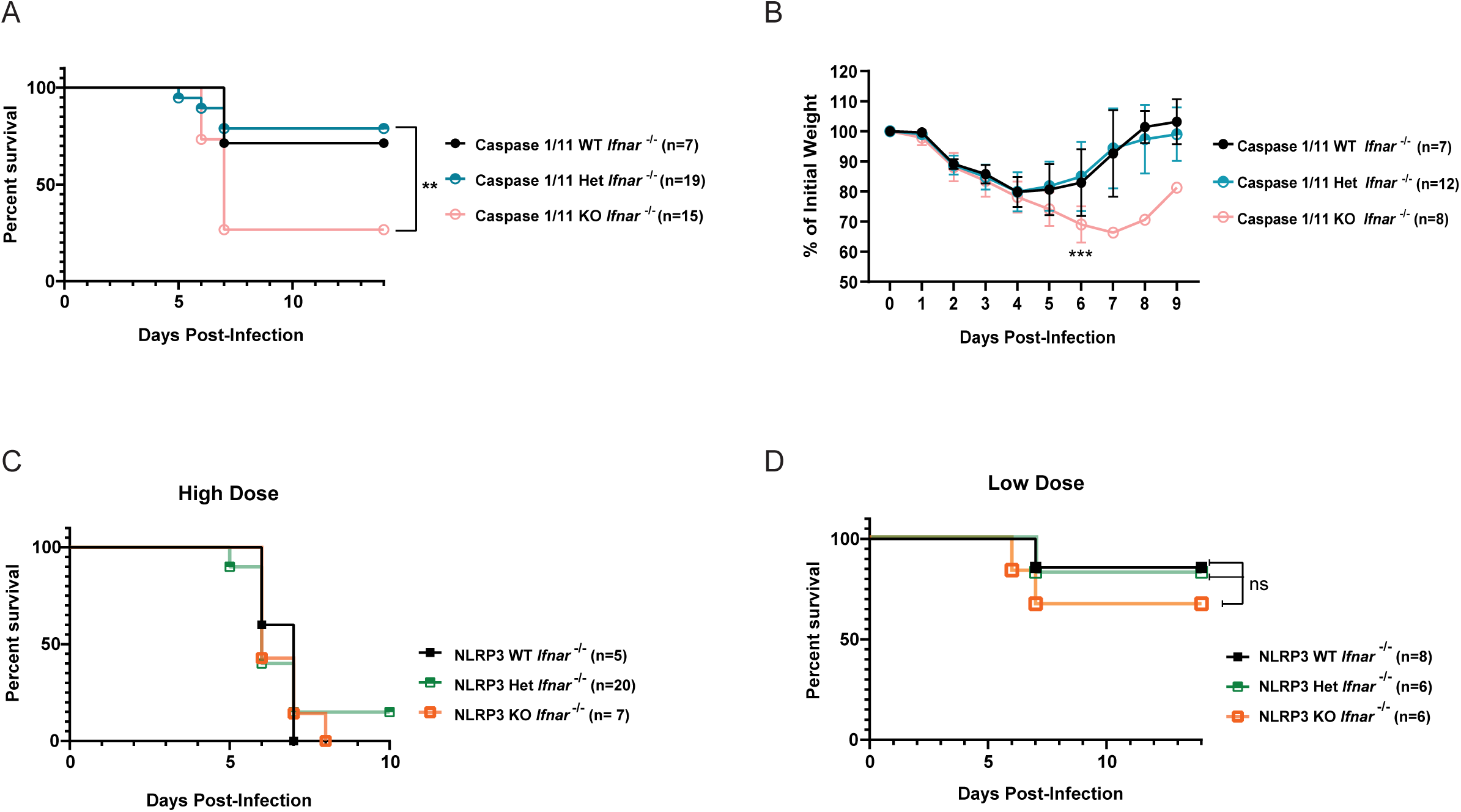
The inflammasome is protective during DENV infection. **(A-B)** Survival curves **(A)** and weight loss over time **(B)** of *Casp1/11^+/+^Ifnar ^-/-^* (Caspase 1/11 WT), *Casp1/11 ^+/-^Ifnar ^-/-^* (Caspase 1/11 Het), or *Casp1/11 ^-/-^Ifnar ^-/-^* (Caspase 1/11 KO) littermates infected intravenously with 3 x 10^5^ PFU of DENV2 D220. Survival was monitored over 14 days. Weight loss was monitored over 9 days. Numbers in parentheses indicate the numbers of mice in each group. **p<0.01. Statistical significance was determined by Mantel– Cox log-rank test **(A)** or two-way ANOVA with Holm-Sidak’s multiple comparisons test **(B)**. **(C-D)** Survival curves of *Nlrp3 ^+/+^Ifnar ^-/-^* (NLRP3 WT)*, Nlrp3 ^+/-^Ifnar ^-/-^* (NLRP3 Het), or *Nlrp3^-/-^Ifnar ^-/-^* (NLRP3 KO) littermates infected intravenously with a 5 x 10^5^ PFU (High Dose) **(C)** or 7.5 x 10^4^ PFU (Low Dose) **(D)** of DENV2 D220 and monitored over 10 days. Numbers in parentheses indicate the numbers of mice in each group.

## Discussion

In this study, we demonstrate that the inflammasome pathway is activated by DENV NS1 in mouse BMDMs and human THP-1 macrophages. Interestingly, we find that DENV NS1-induced inflammasome activation is independent of the NLRP3, AIM2 and Pyrin inflammasomes in BMDMs and that NLRP3 deficiency in mice does not affect the outcome of DENV infection, in contrast to previous reports ascribing a pathogenic role to the NLRP3 inflammasome during DENV infection.^49, 52^ Instead, we find that inflammasome activation may play a protective role during DENV infection, as caspase-1/11-deficient mice are more susceptible to DENV infection compared to their caspsase-1/11-functional littermates.

Our study experimentally assessed the contribution of inflammasome activation to DENV infection *in vivo* using genetically deficient mice, in contrast to previous studies. In one study, investigators sought to define the contribution of DENV M to DENV pathogenesis and found that NLRP3-deficient mice infected with an adeno-associated virus expressing DENV M showed less pathology than WT controls.^49^ However, it is not clear how generalizable this model is to DENV pathogenesis, as the study relied on expression of DENV M protein by an adeno-associated virus rather than by infection with DENV. Our study differs in that we use a DENV2 strain (D220) in a well-characterized IFNAR-deficient mouse model, which has been previously shown to more readily model features of DENV pathogenesis observed in severe disease in humans, such as vascular leak.^64, 65^ Using this model, we found that caspase-1/11-deficient mice are more susceptible to DENV infection than their caspase-1/11 functional littermates, implicating the inflammasome pathway as a protective pathway during DENV infection. Further, we find that NLRP3-deficient, IFNAR-deficient mice are not protected from lethal DENV infection and succumb to infection at similar rates as NLRP3-sufficient IFNAR-deficient mice, thus suggesting that the NLRP3 inflammasome does not contribute to pathogenesis in the context of DENV infection, at least in our DENV D220 mouse model of vascular leak syndrome. Another study showed that treatment of mice with IL-1 receptor antagonist can reduce pathology during DENV infection in IFNAR-deficient mice.^53^ Our results do not necessarily conflict with this prior study since there can be IL-1β-independent effects of inflammasome activation.^66^ Further studies are needed to understand the mechanistic basis of caspase-1/11-mediated protection during DENV infection.

The timing and magnitude of inflammasome activation are key factors in determining whether activation is protective or detrimental to the host. Administration of the NLRP3 inhibitor MCC950 at days 1 and 3 after influenza A virus (IAV) infection led to hyper-susceptibility to lethality, whereas treatment on days 3 and 5 post-infection protected mice against IAV-induced disease.^48^ Thus, in this context, the NLRP3 inflammasome plays both a protective role early in infection and a pathogenic role later in infection. The use of genetic models in our study implicates a protective role for inflammasome activation early in DENV infection as well, though our data does not preclude the possibility of inflammasome activation being pathogenic in later stages of infection.

In addition, increased inflammasome activation was observed under antibody-dependent enhancement (ADE) conditions during a secondary DENV infection.^67, 68^ ADE is a phenomenon whereby cross-reactive non-neutralizing anti-DENV antibodies elicited from a primary DENV infection facilitate Fcγ receptor-mediated viral entry and replication in immune cells during a secondary DENV infection with a different serotype, resulting in higher viremia and increased immune activation.^69, 70^ Thus, the altered viremic and immune context of ADE may also determine whether inflammasome activation still plays a protective role or contributes to DENV disease. While our study ascribes a protective role for inflammasome activation during DENV infection, more investigation is needed to understand whether targeting the inflammasome pathway at specific times and under ADE conditions can lead to therapeutic benefit during severe dengue.

Our data raises some interesting questions regarding the mechanism of DENV NS1-induced inflammasome activation. While we establish that DENV NS1 induces inflammasome activation independently of the NLRP3 inflammasome using multiple orthogonal genetic and chemical approaches, the identity of the inflammasome activated by DENV NS1 is unknown. CRISPR-Cas9-mediated knockout of other, well-studied inflammasomes such as Pyrin and AIM2 had no effect on DENV NS1-induced inflammasome activation, suggesting that these inflammasomes may not be involved or may be redundant with each other. Further work is needed to establish whether DENV NS1 may activate other inflammasomes such as NLRP1B, CARD8, NLRP6 and NLRP10 inflammasomes, as well as to provide deeper insight into the mechanisms behind inflammasome sensing of DENV NS1 in macrophages.^71, 72^

Interestingly, we find here that NS1 can activate the inflammasome pathway through a CD14-dependent pathway without triggering detectable cell death. It has become increasingly appreciated that inflammasome activation and the formation of GSDMD pores in cell membranes do not necessarily lead to pyroptotic cell death, but it is unknown how different inflammasome stimuli induce different cell fates^39^. Previous studies have implicated CD14 as a receptor for oxidized phospholipids such as 1-palmitoyl-2-glutaryl-sn-glycero-3-phosphocholine (PGPC) and 1-palmitoyl-2-(5 -oxo-valeroyl)-sn-glycero-3-phosphocholine (POV-PC); these phospholipids engage CD14 to promote the release of IL-1β from living macrophages via a pathway dependent on caspase-11, caspase-1 and NLRP3.^40, 63, 73^ We found that, similar to POV-PC and PGPC, DENV NS1 can deplete CD14 from the surface of macrophages and is dependent on CD14 to activate the inflammasome pathway. Since LPS is a canonical ligand of CD14 and cytosolic LPS can activate the non-canonical inflammasome pathway, it was critical to eliminate the potential role of LPS in our studies. We primed cells with PAM_3_CSK_4_ and regularly tested DENV NS1 stocks for LPS contamination to ensure that LPS exposure was not driving IL-1β release in our experiments. Importantly, non-canonical inflammasome activation by cytosolic LPS is dependent on the NLRP3 inflammasome, whereas our studies indicate that DENV NS1 activates an inflammasome pathway that is independent of NLRP3. Thus, it is likely that DENV NS1-induced inflammasome activation is not due to contaminating LPS; rather, we propose that, like oxidized phospholipids, DENV NS1 utilizes CD14 to induce IL-1β release in macrophages. However, the mechanism by which oxidized phospholipids enhance IL-1β release remains poorly understood, and further work will be required to determine whether DENV NS1 operates via a similar or distinct mechanism.

A previous study reported that NS1-associated lipid cargo is enriched in triglycerides, cholesterol, and phospholipids, lipids commonly found in cell membranes.^11^ We speculate that NS1 could act as a carrier of oxidized phospholipids generated from infected cells and subsequently be detected by macrophages at sites distal from infection, activating an inflammasome and cytokine response. We were unable to determine whether oxidized phospholipids were present within the lipid cargo of our NS1 in this study; however, it would be interesting to explore whether the inflammatory capacity of NS1 is ultimately modulated by the lipids within the lipid cargo.

Overall, our results provide insight into interactions between DENV NS1 and macrophages and the role of NS1 in protection. Our current study suggests that NS1-myeloid cell interactions can be protective during DENV infection and that activation of pro-inflammatory circuits during DENV infection can be beneficial. Thus, we find that the activation of pro-inflammatory immune responses does not always lead to detrimental outcomes during DENV infection, contrary to many studies in the field.^22, 27, 74^ Instead, a more nuanced view accounting for the timing and magnitude of the inflammatory response may be a crucial aspect of understanding both the beneficial and detrimental aspects of inflammation in DENV infection. Our data suggest that further investigation into understanding the delicate balance and precise contexts in which cytokines can be protective and/or pathogenesis during DENV infection will be crucial for developing novel therapeutics and identifying the best biomarkers to assess risk of progression to severe disease.

## Acknowledgements

We thank Dr. Patrick Mitchell at the University of Washington and Elizabeth Turcotte from Dr. Russell Vance’s laboratory for provision of key reagents and helpful discussions about inflammasome biology. Additionally, we thank Kristen Witt, Dr. Sarah Stanley, and Marietta Ravesloot-Chavez for training on CRISPR-Cas9 genetic editing in macrophages and access to their nucleofectors. We also thank Xinyi Feng, Elin Lee, Eduarda Lopes, Pedro Henrique Nascimento Carneiro da Saliva for assistance with animal colony maintenance and experiments; Samantha Hernandez, Maria Jose Andrade, Marco Chapa, and Claudia Sanchez San Martin for administrative and lab management support; and Nicholas Tzuning Lo, Elias Duarte, Sandra Bos, and P. Robert Beatty for their many helpful discussions. This study was supported by NIH grants R01 AI124493 and R01 AI168003 to E.H. R.E.V. is an Investigator of the Howard Hughes Medical Institute. S.B.B was partially supported as an Open Philanthropy Awardee of the Life Science Research Foundation.

## Author Contributions

Conceptualization: M.P.W., S.B.B., R.E.V., and E.H.; Methodology: M.P.W., E.Y.W.J., S.B.B., and E.H.; Formal Analysis: M.P.W.; Investigation: M.P.W., E.Y.W.J., S.C.C., P.W., F.P., B.C.R., S.F.B., and L.M.D.; Resources: L.M.D., F.T.G.S., L.S.V.B., C.M.R., R.E.V., and E.H.; Visualization: M.P.W.; Writing-Original Draft: M.P.W., S.B.B., and E.H., Writing-Review and Editing: M.P.W., E,Y.W.J., S.C.C., F.P., L.M.D., F.T.G.S., C.M.R., S.B.B, R.E.V, and E.H.; Supervision: S.B.B, R.E.V, and E.H.; Funding Acquisition: E.H.

## Materials and Methods

### Mice

All mice were bred in-house in compliance with Federal and University regulations. All experiments involving animals were pre-approved by the Animal Care and Use Committee (ACUC) of UC Berkeley under protocol AUP-2014-08-6638-2 and maintained under specific pathogen-free conditions. Wildtype (WT) C57BL/6 mice were obtained from Jackson Labs and used for preparation of bone marrow-derived macrophage (BMDMs) . *Nlrp3*^-/-^ and *Casp-1/11*^-/-^ C57BL/6 mice were provided by Dr. Russell Vance and maintained in the Harris lab mouse colony. *Nlrp3*^+/-^ and *Casp 1/11*^+/-^ mice were bred to generate the corresponding WT ^(+/+^) or deficient (^-/-^) litermates mice the corresponding ^(+/+^) or deficient (^-/-^) litermates mice for all experiments. For *in vivo* experiments involving DENV2 infection, *Nlrp3*^-/-^ and *Casp 1/11*^-/-^ were bred with C57BL/6 mice deficient in the interferon α/β receptor (*Ifnar*^-/-^) to generate doubly-deficient *Nlrp3^+^*^/-^ x *Ifnar*^-/-^ or *Casp-1/11*^+/-^ x *Ifnar*^-/-^ mice by backcrossing *Nlrp3*^-/-^ and *Casp-1/11*^-/-^ mice with *Ifnar*^-/-^ mice over 6 generations. The genotypes of mice used for breeding were tracked via PCR and gel electrophoresis. *Nlrp3^+^*^/-^ x *Ifnar*^-/-^ or *Casp-1/11*^+/-^ x *Ifnar*^-/-^ mice were bred to generate littermate WT, heterozygous, or KO mice at the *Nlrp3*^-/-^ or *Casp-1/11*^-/-^ allele.

Individual PCR reactions were utilized to confirm the WT genotype and genetically deficient genotype for *Casp-1/11* and *Nlrp3*. The primers for each of these reactions were as follows: *Caspase 1/11* WT (*CATGCCTGAATAATGATCACC* and *GAAGAGATGTTACAGAAGCC*), *Caspase 1/11*-deficient(*GCGCCTCCCCTACCCGG* and *CTGTGGTGACTAACCGATAA*), *Nlrp3* WT (*CCACCTGTCTTTCTCTCTCTGGGC* and *CCTAAGGTAAGCTTTTGTCACCCAGG*), *Nlrp3*-deficient (*TTCCATTACAGTCACTCCAGATGT* and *TGCCTGCTCTTTACTGAAGG*). To determine the IFNAR genotype of the mice, a multiplex PCR protocol consisting of three primers was used (*CGAGGCGAAGTGGTTAAAAG*, *ACGGATCAACCTCATTCCAC*, and *AATTCGCCAATGACAAGACG*).

### Viral stocks and proteins

A mouse-attenuated strain of DENV2, D220, was utilized for infection of *Ifnar*^-/-^ mice.^64^ The D220 strain was derived from the Taiwanese DENV2DENV2 isolate PL046 and is a further modification of the D2S10 strain, as described previously.^64^ Viral stocks were titered using focus-forming assays on Vero cells. All virus stocks were confirmed to be mycoplasma-free.

Recombinant DENV2 NS1 (Thailand/16681/84) was produced in mammalian HEK293 cells or purchased from The Native Antigen Company (Oxford, UK). All NS1 stocks were certified to be endotoxin-free and >95% purity.

### DENV mouse model

Six-to eight-week-old littermate mice of either gender were challenged with DENV via retroorbital intravenous (IV) injection of the indicated plaque-forming units (PFU) of the DENV2 D220 strain. Mice were observed for morbidity and mortality over a 2-week period. Morbidity of mice was assessed utilizing a standardized 1-5 scoring system as follows: 1 = no signs of lethargy, mice are considered healthy; 2 = mild signs of lethargy and fur ruffling; 3 = hunched posture, further fur ruffling, failure to groom, and intermediate level of lethargy; 4 = hunched posture with severe lethargy and limited mobility, while still being able to cross the cage upon stimulation; and 5 = moribund with limited to no mobility and inability to reach food or water.(REF) Mice scored as 4 were monitored twice per day until recovery or until reaching moribund status. Moribund mice were euthanized immediately. Mice were also weighed to measure weight changes throughout the infection period.

### BMDM generation

BMDMs were generated from 8-13-week-old WT C57BL/6 mice purchased from Jackson Laboratories or 8-13-week-old C57BL/6 littermate mice of the following genotypes: *Casp-1/11*^-/-^, *Casp-1/11*^+/+^, *Nlrp3*^-/-^, and *NLRP3*^+/+^. Additionally, BMDMs were generated from 8-13-week-old WT C57BL/6 mice purchased from Jackson Laboratories.*Nlrp3*^+/+^. Bone marrow was extracted from the femur and tibia of dissected mice and plated on non-tissue culture-treated 15-cm Petri dishes at a density of 1 x 10^7^ cells per plate in macrophage differentiation medium (DMEM, Gibco) supplemented with 2mM L-glutamine, 10% fetal bovine serum (FBS; Corning), 1% penicillin/streptomycin (Pen/Strep, Gibco), and 10% macrophage colony-stimulating factor (M-CSF) containing supernatant solution harvested from 3T3-CSF cells and cultured at 37°C with 5% CO_2_ for 7 days. On day 3 of incubation, cells were supplemented with additional macrophage differentiation medium. On day 7, differentiated BMDM cells were harvested by incubating the plated cells in phosphate-buffered saline without calcium and magnesium (PBS) at 4°C for 20 minutes (min). The BMDM cells were then removed from the plate by gentle spraying with PBS, resuspended in DMEM supplemented with 1% Pen-Strep, 40% FBS, and 10% DMSO and frozen in liquid nitrogen until future use.

### BMDM inflammasome activation assay

A macrophage-based assay was adapted from previously described protocols to assess inflammasome activation in BMDMs.^60^ Frozen BMDMs were thawed and plated in 24-well or 96-well tissue culture-treated, flat-bottom plates in complete DMEM (DMEM + 2mM L-glutamine + 10% FBS + 1% Pen/Strep) at a density of 1x10^6^ or 2x10^5^ cells per well, respectively. After plating, cells were left to rest at 37°C and 5% CO_2_ overnight. BMDMs were then primed using 1μg/mL Pam_3_CSK_4_ (InvivoGen) for 17 hours (h) or left untreated. Primed BMDMs were stimulated with DENV NS1, 5μM nigericin (Invivogen), or left untreated. After 24h, supernatants were spun down at 10,600 x g for 10 minutes, and the cell-free supernatant was collected. BMDM layers were washed twice with PBS and then lysed in RIPA buffer (150mM NaCl, 1% Nonidet P-40, 0.5% sodium deoxycholate, 0.1% sodium dodecyl sulfate (SDS), 50mM Tris pH 7.4) supplemented with protease inhibitor (Roche). Supernatants and cell lysates were stored at -80°C until further analysis. For experiments involving inhibitors, the NLRP3 inhibitor MCC950 (InvivoGen) or caspase-1 inhibitor Ac-YVAD-cmk (InvivoGen) were added at indicated concentrations 30 min prior to treatment with DENV2 NS1 or nigericin.

### Cell culture

HEK-Blue-IL-1β cells were obtained from Invivogen (catalog # hkb-il1b) and grown in complete medium containing DMEM, 10% FBS, 1% Pen/Strep, 100μg/mL Zeocin (Invivogen), and 200μg/mL Hygromycin B Gold (Inviviogen). WT THP-1 cells or *casp-1^-/-^* THP-1 were purchased from Invivogen and grown in complete medium containing RPMI (Gibco), 10% FBS, and 1% Pen/Strep. HEK-Blue IL-1β reporter cells were grown and assayed in 96-well plates. All cell lines were routinely tested for mycoplasma by PCR kit (ATCC, Manassas, VA).

### THP-1 inflammasome activation assay

To assess inflammasome activation in THP-1 macrophages, WT or *casp-1^-/-^*THP-1 human monocytes were differentiated into macrophages in 10ng/mL phorbol 12-myristate 13-acetate (PMA) and primed with medium or 1μg/mL LPS for 4h. Primed macrophages were treated with 10μg/mL DENV NS1 or left untreated (LPS only). 18h later, supernatants were collected. Cells were stimulated with 5μM nigericin for 2h as a positive control. Supernatants were then assessed for bioactive IL-1β using a HEK-Blue IL-1β reporter assay.

To assess cleavage of caspase-1 in THP-1 macrophages, WT THP-1 human monocytes were differentiated into macrophages in 10ng/mL PMA and primed with medium or 100 ng/mL Pam_3_CSK_4_ for 17h. Primed macrophages were then treated with DENV NS1 or 5μM nigericin in RPMI + 2% FBS + 1% Pen/Strep for 24h. Supernatants were collected, and cells were lysed with RIPA buffer. Proteins in supernatant were precipitated by methanol/chloroform precipitation and resuspended in 50uL of 1X SDS-PAGE sample buffer (0.1% β-mercaptoethanol, 0.0005% bromophenol blue, 10% glycerol, 2% SDS, 63mM Tris-HCl pH 6.8).

### HEK-Blue IL-1β reporter assay

To quantify the levels of bioactive IL-1β released from cells, we employed HEK-Blue IL-1β reporter cells. In these cells, binding of IL-1β to the surface receptor IL-1R1 results in the downstream activation of NF-κB and subsequent production of secreted embryonic alkaline phosphatase (SEAP) in a dose-dependent manner. SEAP levels were detected using a colorimetric substrate assay, QUANTI-Blue (Invivogen) by measuring an increase in absorbance at OD655. Culture supernatant from THP-1 cells was added to HEK-Blue IL-1β reporter cells plated in 96-well format in a total volume of 200 μl per well. After 24h, SEAP levels were assayed by adding 20 μl of the supernatant from HEK-Blue IL-1β reporter cells to 180 μl of QUANTI-Blue colorimetric substrate following the manufacturer’s protocol. After incubation at 37°C for 30-60 min, absorbance at OD_655_ was measured on a microplate reader.

### CRISPR-Cas9-mediated gene editing in primary BMDMs

CRISPR-Cas9 gene editing was performed in WT BMDMs to knock out specified genes using a nucleofection-based approach.^75^ Bone marrow from WT B6 mice was isolated, and cells were differentiated as previously described. On day 5 of macrophage differentiation, BMDMs were harvested and resuspended in P3 Primary Cell Nucleofector Solution with Supplement (Lonza). Separately, Cas9-ribonuclear protein (RNP) complexes were made by incubating *S. pyogenes* Cas9 with 2 nuclear localization signals (SpCas9-2NLS, Synthego) and 2 guide RNAs per gene (Synthego), together with Alt-R^TM^ Cas9 Electroporation Enhancer (IDT) at room temperature for 25 min. BMDMs (4x10^6^ per gene target) were mixed with RNP complex, and cells were nucleofected in 16-well Nucleocuvette strips (Lonza) using a 4D-Nucelofector^®^ (Lonza).

Nucleofected cells were then seeded in non-tissue culture-treated 15-cm Petri dishes in macrophage differentiation medium and differentiated for an additional 5 days, with medium changes every 2 days. After differentiation, gene-edited macrophages were harvested and used in the previously described inflammasome activation assay. Gene-edited macrophages were stimulated with inflammasome activators for 48h before supernatants were assessed for IL-1β.

Additionally, primed, gene-edited macrophages were lysed in RIPA buffer, and cell lysates were frozen for validation of efficient gene knockout by Western blot.

### Cytokine and lactase dehydrogenase (LDH) quantification

Cytokine levels in cell supernatants were assessed using the mouse IL-1β/IL-1F2 DuoSet ELISA kit (R&D Systems) and mouse TNF-α DuoSet ELISA kit (R&D Systems) according to the manufacturer’s instructions. ELISA plates were measured at OD_450_ using a microplate reader, and cytokine levels were quantified by interpolation using a standard curve. Supernatants were assayed for LDH release immediately after stimulation time courses per the manufacturer’s protocol from the CytoTox 96^®^ Non-Radioactive Cytotoxicity Assay (Promega). Measurements of absorbance readings were performed on a microplate reader at wavelengths of 490 nm and 680 nm. LDH release was assessed as a percentage of background-subtracted maximum LDH values from lysed cells.

### SDS-PAGE and Western blot

Cell supernatants and lysates were diluted in 6X SDS-PAGE protein sample buffer (360mM Tris pH 6.8, 12% SDS, 18% β-mercaptoethanol, 60% glycerol, 0.015% bromophenol blue), boiled for 10 min at 95°C, and resolved using SDS-PAGE. The proteins were then transferred onto a nitrocellulose membrane, washed 3 times with Tris-buffered saline with 0.1% Tween20 (TBS-T) and probed with primary antibodies diluted in 5% non-fat dry milk in TBS-T at 4°C overnight.

The membrane was then washed 3 times with TBS-T and probed with secondary antibodies in 5% non-fat dry milk in TBS-T rocking at room temperature for 2 hours. The membrane was then washed 3 times with TBS-T and 2 times in TBS and developed using SuperSignal West Pico PLUS Chemiluminescence reagent (ThermoFisher). The resulting membrane was imaged on a BioRad ChemiDoc system and visualized using Image Lab software (BioRad). The antibodies and working dilutions used were as follows: goat-α-mouse IL-1β, 1:1000 (R&D Systems, AF-401-NA); rabbit-α-NLRP3, 1:1000 (Cell Signaling, D4D8T); rabbit-α-mouse AIM2, 1:500 (Cell Signaling, 63660), α-mouse Caspase-1 (p20) 1:1000 (Adipogen, Casper-1), α-human Caspase-1 (p20), 1:1000 (Adipogen, Bally-1); α-Pyrin, 1:1000 (Abcam, EPR18676); goat-α-mouse IgG HRP, 1:5000 (Biolegend, 405306); donkey-α-rabbit IgG HRP, 1:5000 (Biolegend, 405306); α-actin HRP, 1:2000 (Santa Cruz Biotechnologies, sc-8432).

### Flow Cytometry

BMDMs were primed with 1μg/mL Pam_3_CSK_4_ for 17h, then stimulated with 5μg/mL *E. coli* LPS, 10μg/mL DENV NS1, or 5μM nigericin for the indicated time periods at 37°C. Cells were washed twice with PBS, then incubated in PBS at 4°C for 10 min and scraped to suspend cells. Suspended cells were stained with either Live/Dead Fixable Far Red Stain (ThermoFisher) or APC-α-mouse CD14 primary antibody (clone Sa2-8; Thermo Scientific) on ice in the dark for 30 minutes. Purified rat α-mouse CD16/CD32 (Mouse FcBlock; Becton Dickenson) was used as the blocking reagent to reduce non-specific binding of the antibodies. The stained cells were then washed twice with 1ml cold FACS buffer (1% Bovine Serum Albumin [Sigma] and 1% Purified Mouse IgG [Invitrogen] in PBS) and fixed in 500μL of 4% paraformaldehyde at room temperature. Cells were washed once in PBS and kept in 500μL PBS at 4°C in the dark until analysis with an Intellicyt iQue3 Screener (Sartorius). For viability analysis, a dead-cell gate was set based on unstained cell controls, and the percentage of singlet cells in the dead-cell gate compared to all singlet cells was calculated. For CD14 expression, the mean fluorescence intensity (MFI) of CD14 from unstimulated or stimulated cells was recorded. The percentage of surface receptor staining at 30 min, which is the ratio of the MFI values measured from the stimulated cells to those measured from the unstimulated cells, was plotted to reflect the efficiency of receptor endocytosis. At least 10,000 events were acquired per sample for analysis.

### Statistics

All quantitative analyses were conducted and all data were plotted using GraphPad Prism 8 Software. Data with error bars are represented as mean ± SEM. Statistical significance for experiments with more than two groups was tested with two-way ANOVA with multiple comparison test correction as indicated.

